# Phylogenetic tree shapes resolve disease transmission patterns

**DOI:** 10.1101/003194

**Authors:** Caroline Colijn, Jennifer Gardy

## Abstract

**Background and Objectives:** Whole genome sequencing is becoming popular as a tool for understanding outbreaks of communicable diseases, with phylogenetic trees being used to identify individual transmission events or to characterize outbreak-level overall transmission dynamics. Existing methods to infer transmission dynamics from sequence data rely on well-characterised infectious periods, epidemiological and clinical meta-data which may not always be available, and typically require computationally intensive analysis focusing on the branch lengths in phylogenetic trees. We sought to determine whether the topological structures of phylogenetic trees contain signatures of the transmission patterns underlying an outbreak.

**Methodology:** We use simulated outbreaks to train and then test computational classifiers. We test the method on data from two real-world outbreaks.

**Results:** We show that different transmission patterns result in quantitatively different phylogenetic tree shapes. We describe topological features that summarize a phylogeny’s structure and find that computational classifiers based on these are capable of predicting an outbreak’s transmission dynamics. The method is robust to variations in the transmission parameters and network types, and recapitulates known epidemiology of previously characterized real-world outbreaks.

**Conclusions and implications:** There are simple structural properties of phylogenetic trees which, when combined, can distinguish communicable disease outbreaks with a super-spreader, homogeneous transmission, and chains of transmission. This is possible using genome data alone, and can be done during an outbreak. We discuss the implications for management of outbreaks.

## 1 Introduction

Whole-genome sequence data contain rich information about a pathogen population from which several evolutionary parameters and events of interest can be inferred. When the population in question comprises pathogen isolates drawn from an outbreak or epidemic of an infectious disease, these inferences may be of epidemiological importance, able to provide actionable insights into disease transmission. Indeed, since 2010, several groups have demonstrated the utility of genome data for revealing pathogen transmission dynamics and identifying individual transmission events in outbreaks [Stadler et al., 2012, Köser et al., 2012, Walker et al., 2012, Grad et al., 2012, Török et al., 2013, Kato-Maeda et al., 2013, Ypma et al., 2013, Jombart et al., 2014, Didelot et al., 2014], with the resulting data now being used to inform public health’s outbreak management and prevention strategies. To date, these reconstructions have relied heavily on interpreting genomic data in the context of available epidemiological data, drawing conclusions about transmission events only when they are supported by both sequence data and plausible epidemiological linkages collected through field investigation and patient interviews.

Given the rapidly growing interest in this new field of genomic epidemiology, several recent studies have explored whether transmission events and patterns can be deduced from genomic data alone. Phylogenies derived from whole-genome sequence data can be compared to theoretical models describing how a tree should look under particular processes; this has been done for viral sequence data over the past several decades [Pybus et al., 2001, Pybus and Rambaut, 2009]. For example, predicted branch lengths from sequences modeled using birth-death processes can be compared to branch lengths in trees inferred from viral sequence data to explore transmission patterns [Stadler and Bonhoeffer, 2013, Stadler et al., 2012, Leventhal et al., 2012, Stadler, 2011]. The field of linking properties of pathogen phylogenies to underlying dynamics is termed *phylodynamics*, coined by Grenfell et al [Grenfell et al., 2004]. Tools from coalescent theory have been adapted to pathogen transmission; where coalescent theory describes probability distributions on trees under a given model for the population size, epidemiological versions take into account the relationship between pathogen prevalence (population size) as well as incidence [Volz, 2012, Volz et al., 2012]. These approaches are powerful, but are computationally intensive and have not explicitly focused on another potential source of information within a phylogeny - *tree shape*.

The number of different phylogenetic tree shapes on *n* leaves is a combinatorially exploding function of *n* (there are (2*n−*3)(2*n−*5)(2*n−*7)*…*(5)(3)(1) rooted labelled phylogenetic trees, or approximately 10^184^ trees on 100 tips, compared to approximately 10^80^ atoms in the universe). For the increasingly large outbreak genome datasets being obtained and analysed (390 [Walker et al., 2012], 616 [Croucher et al., 2013] and recently 1000 [Casali et al., 2014] bacterial genomes), the numbers of possible tree shapes are effectively infinite. In the homogeneous birth (Yule) model, the distribution of labelled histories (tree shape together with the ordering of internal nodes in time) is uniform, so that there is a close relationship between the branching times and the tree shapes [Rosenberg, 2006]. Perhaps for this reason, tree shapes have not typically been seen as very informative. However, for bacterial pathogens, particularly those with long durations of carriage and variable infectious rates, there is variability in the infection process which is not captured by homogeneous models. This motivates asking the question: does tree shape carry epidemiological information? Recent work indicates that tree shape reveals aspects of the evolution of viral pathogens [Norström et al., 2012, Leventhal et al., 2012, Robinson et al., 2013, Poon et al., 2013, Frost and Volz, 2013], but to date we do not have methods to exploit tree shape in an analysis of pathogen transmission dynamics, built upon simulated data and validated using real-world outbreak data.

Host contact network structure is one of the most profound influences on the dynamics of an outbreak or epidemic, and outbreak management and control strategies depend heavily upon the type of transmission patterns driving an outbreak. It is reasonable to expect that pathogen genomes spreading over different contact network structures - chains, homogenous networks, or networks containing super-spreaders, as illustrated in Figure 1 - would accrue mutations in different patterns, leading to observably different phylogenetic tree shapes. We therefore characterized the structural features of phylogenetic trees arising from the simulated evolution of a bacterial genome as it spreads over multiple types of contact network. We found simple topological properties of phylogenetic trees that, when combined, can be used to classify trees according to whether the underlying process is chain-like, homogenous, or super-spreading, demonstrating that phylogenetic tree structure can reveal transmission dynamics. We use these properties as the basis for a computational classifer, which we then use to classify real-world outbreaks. We find that the computational predictions of each outbreak’s overall transmission dynamics are consistent with known epidemiology.

**Figure 1:**
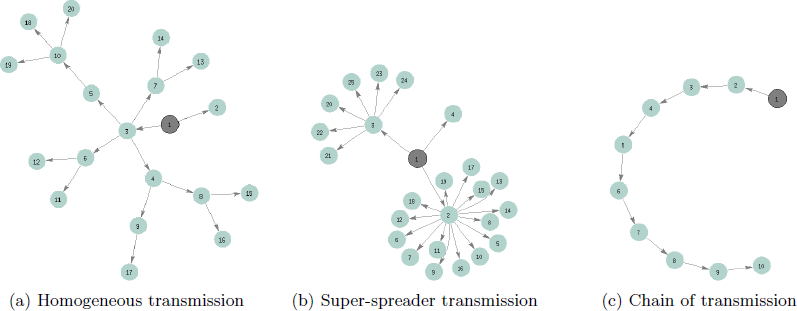
Schematic illustration of different kinds of transmission networks. The index case is marked in grey.

## 2 Materials and Methods

### Transmission model

We simulated disease transmission networks with three different underlying transmission patterns: homogeneous transmission, transmission with a super-spreader, and chains of transmission. Each simulation started with a single infectious host who infects a random number of secondary cases over his or her infectious period; each secondary case infects others, and so on, until the desired maximum number of cases is reached. The models share two key parameters: a transmission rate *β* and a duration of infection parameter *D*. Our baseline values are *β* = 0.43 per month and *D* = 3 months, reflecting a basic reproduction number of 1.3. This is also the mean number of secondary infections for each infectious case. We do not consider depletion of susceptible contacts over time (saturation) as we model small growing outbreaks at or near the beginning of their spread in a community, and our data (for TB in a developed setting) suit this assumption.

The homogeneous transmission model assigns each infectious host a number of secondary infections drawn from a Poisson distribution with parameter *R*_0_ = *βD*. New infections are seeded uniformly in time over the host’s infectious period. In the super-spreader model, one host (at random in the first 5 hosts) seeds 7-24 new infections (uniformly at random), and all other hosts are as in the homogeneous transmission model. In the chain-of-transmission model, almost all hosts infect precisely one other individual. However, 2 (with probability 2/3) or 3 (with probability 1/3) of the hosts infect two other individuals, so that the transmission tree consists of several chains of transmission randomly joined together.

Durations of infection are drawn from a Γ distribution with a shape parameter of 1.5 and a scale parameter of *D/*1.5. To reflect transmission of a chronically-infecting pathogen, such as *Mycobacterium tuberculosis*, cases were infectious for between 2 and 14 months with an average specified by The mean infectious period was 4.3 months; a histogram is shown in Figure S2. We simulated 1000 outbreaks containing a super-spreader, 1000 with homogeneous transmission and 1000 chain-like outbreaks. These used a fixed parameter set; we also performed a sensitivity analysis using alternative parameters. To ensure that the size of the outbreak did not affect the tree shape and classification, we simulated outbreaks with 32 hosts - a similar size as the real-world outbreaks we later investigated. We consider the effects of phylogenetic noise in the Supplementary Material.

### Genealogies and phylogenies from the process

We extracted the true genealogical relationships as a full rooted binary tree (a “phylogeny”), with tips corresponding to hosts and internal nodes corresponding to transmission events among the hosts, as follows. The outbreak simulations create lists of who infected whom and at what time. Each host also has a recovery time. We sort the times of all of the infection events, and proceed in reverse order. The last infection event must correspond to a “cherry”, ie it must have two tip descendants, one corresponding to the infecting host and one to the infectee. For all other infection events proceeding in reverse order through the transmission, we create an internal node, and determine its descendants by determining whether the infector and the infectee went on to infect anyone else subsequently. If not, then the node’s descendants are the infector and infectee at the time of sampling. If so, then the descendant represents the infector or infectee at the time of their next transmission. The tree is rooted at the first infection event. Branch lengths correspond to the times between infection events or, for tips, the time between the infection event and the time of sampling. This approach uses the simplifying assumption that branching points in the pathogen genealogy correspond to transmission events, as is done in almost all phylodynamic methods (see [Stadler et al., 2012, Volz et al., 2012, Frost and Volz, 2013]). However, where there is in-host pathogen diversity transmission events do not correspond to phylogenetic branching points [Didelot et al., 2014, Ypma et al., 2013]. We comment on the constraints tree shape places on the space of possible transmission trees consistent with a phylogeny in [Didelot et al., 2014].

In the main text, we use the true genealogical relationships among the hosts in our outbreak, extracted from the simulations - this reduces phylogenetic noise and it allows us to compare the resulting trees to 1000 samples of the BEAST posterior timed phylogenies derived from WGS data from the two real-world outbreaks. To determine how sensitive our approach is to phylogenetic noise, we also classified the outbreaks using neighbour-joining phylogenies derived from simulated gene sequences (Supplementary Information).

### Topological summaries of trees

Eleven summary metrics were used to summarise the topology of the trees (see Table S1).

1. **Imbalance**. The Colless imbalance [Colless, 1982] is defined as 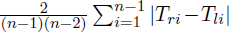 where *n* is the number of tips and *T*_*ri*_ and *T*_*li*_ are the number of tips descending from the left and right sides at internal node *i*. It is a normalised measure of the asymmetry of a rooted full binary tree, with a completely asymmetric tree having imbalance of 1 and a symmetric tree having an imbalance of 0 [Heard, 1992]. The Sackin imbalance [Sackin, 1972] is the average length of the paths from the leaves to the root of the tree.
2. **Ladders, IL nodes** We define the *ladder length* to be the maximum number of connected internal nodes with a single leaf descendant, and we divide it by the number of leaves in the tree. This measure is not unrelated to tree imbalance but is more local - a long ladder motif may occur in a tree that is otherwise quite balanced. For this reason, ladder length may detect trees in which there has been differential lineage splitting in some clades or lineages but where this occurred too locally or in clades that are too small to have affected traditional approaches to characterising rapid expansion in some lineages. Furthermore, traditional ways of detecting positive selection may not be appropriate in this context because the super-spreader, if present, does not pass any advantageous property to descendant infections. The portion of “IL” nodes is the portion of internal nodes with a single leaf descendant.
3. **Maximum width; Maximum width over maximum depth**. The *depth* of a node in a tree is the number of edges between that node and the tree’s root. The *width* of a tree at a depth *d* is defined as the number of nodes with depth *d*. We calculated the maximum width of each tree divided by its maximum depth (max *d*, the maximum depth of any leaf in the tree).
4. **Maximum difference in widths** We compared 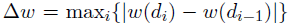 in the trees. This summary reflects the maximum absolute difference in widths from one depth to the next, over all depths *d*_*i*_ in the tree.
5. **Cherries** A cherry configuration is a node with two leaf descendants.
6. **Staircase-ness** We use two measures of the “staircase-ness” of phylogenies defined by Nordstrom et al [Norström et al., 2012]: (1) the portion of sub-trees that are imbalanced (ie that have different numbers of descending tips on the left and right sides) and (2) the average of 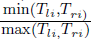 over the internal nodes of the tree.

### Outbreak classification

We trained k-nearest-neighbour (KNN) classifiers using matlab’s Classification KNN.fit function with a Minkowski distance, inverse distance weighting and 100 neighbours. KNN classification was performed on 1000 trees of each type (homogeneous transmission, super-spreaders, chains) using 10-fold cross-validation. The 10 resulting classifiers were then used to classify the groups of simulations in the sensitivity analysis, allowing us to report on the variability of classification results. KNN classification is suitable for sets of data that have any number of groups. Here there were three groups: homogeneous outbreaks, super-spreader outbreaks, and chains of transmission. KNN classifiers’ quality can be assessed with a table reporting how many in each group are correctly classified, and how many are classified into which incorrect group. Alternatively, the quality can be summarised by reporting the portion of each group that is classified correctly.

When there are only two groups to compare, so that classification is binary, better methods are available. One of the most powerful of these is the support vector machine approach. We used a 10-fold cross-validated support vector machine (SVM) to resolve differences between homogeneous transmission vs super-spreader networks. Because SVMs are binary classifiers, their quality can be assessed by reporting the sensitivity (portion of true positives that are classified as positive) and specificity (portion of true negatives that are classed as negatives) of the predictions. The sensitivity and specificity of a classifier trade off with each other, because it is always possible to classify all cases as positive (sensitivity 1 but specificity 0) or all as negative (specificity 1 but sensitivity 0). Classifiers use a cutoff, calling a data point positive if the cutoff is above some threshold, and negative otherwise. The overal quality of a binary classifier can be visualised using a receiver operator characteristic (ROC) curve, which captures the change in sensitivity and specificity of a classifier when its threshold is changed. See [Cristianini and Shawe-Taylor, 2000] for a full discussion of support vector machines and classification.

Here, SVMs were constructed using the svmtrain method in matlab with a linear kernel function. The training data *x*_*i*_ in the *i^’^th* “fold” were the 11 summary metrics for 900 trees derived from each process. The test data were the remaining 100 trees. This was done 10 times (10 “folds” of cross-validation). All training data were from simulations with the baseline set of parameters. The 10 SVMs (one for each “fold”) were tested on the remaining trees using matlab’s svmclassify, which computes

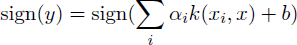

where *α*_*i*_ are weights, *x*_*i*_ are the support vectors, *x* is the input to be classified, *k* is the kernel function and *b* is the bias. These tests were done separately on the different groups of simulated trees. The svmclassify function was modified to return *y* (i.e. the degree to which an outbreak could be considered super-spreading) rather than only the sign of *y* (a binary prediction). We have also performed 10-fold svm classification in R using the e1071 package. Classifiers are available along with a script to profile the structure of a tree in newick format, using the phyloTop package [Boyd and Colijn, 2014] (see Supplementary Information).

### Sensitivity analysis

To determine whether the classifier is robust to different choices of model parameters and to sampling, we simulated three groups of 500 homogeneous and super-spreader outbreaks with (i) randomly selected parameters, (ii) a random sampling density, and (iii) both random parameters and random sampling. Group (i) had randomized parameters in which *β/D* was uniformly distributed between 1.25 and 2.5. Group (ii) had fixed parameters, but the number of cases varied uniformly between 100 and 150, and we sampled only 33 of those cases. The third group had both randomized parameters and random sampling.

To ensure that the classification is detecting variability in the number of secondary cases (ie super-spreading), we performed classification on outbreaks in which we used a negative binomial distribution to determine the numbers of secondary cases in the outbreak. We varied the parameters of the distribution such that the mean number of secondary cases was the same (*R*_0_) but the variance differed, with the expected variance ranging from 2-5 times what it would be in the Poisson (homogeneous transmission) case. We classified the outbreaks using the 10 SVM classifiers obtained under 10-fold cross-validation on the baseline case. We report the mean and standard deviation of the specificity, ie the portion of cases correctly classified as ‘super-spreader’ outbreaks.

To determine whether the classifier is relevant to different *kinds* of models, we applied it to simulated phylogenies described in Robinson et al [Robinson et al., 2013]. In that work, dynamic networks of sexual contacts were created based on random graphs with a Poisson distribution, and with a distribution of contacts derived from the National Survey on Sexual Attitudes and Lifestyles (NATSAL) [Johnson et al., 2001]. See Supplementary material for further details.

### Classification of outbreaks from published genomic data

We used the classifier on phylogenetic trees derived from two real-world tuberculosis outbreak datasets. Outbreak A was previously published [Gardy et al., 2011] and is available in the NCBI Sequence Read Archive under the accession number SRP002589. This dataset comprises 31 *M. tuberculosis* isolates collected in British Columbia over the period 1995-2008 and was sequenced using paired-end 50 bp reads on the Illumina Genome Analyzer II. Outbreak B comprises 33 *M. tuberculosis* isolates collected in British Columbia over the period 2006-2011, and was sequenced using paired-end 75 bp reads on the Illumina HiSeq. The outbreak, sequences and SNPs are presented in [Didelot et al., 2014].

For both datasets, reads were aligned against the reference genome M. tuberculosis CDC1551 (NC002755) using Burrows-Wheeler Aligner (BWA) [Li and Durbin, 2009]. Single nucleotide variants were identified using samtools mpileup [Li et al., 2009] and were filtered to remove any variant positions within 250 bp of each other and any positions for which at least one isolate did not have a genotype quality score of 222. The remaining variants were manually reviewed for accuracy and were used to construct a phylogenetic tree for each outbreak as described above. We apply the classification methods to 1000 samples from the BEAST posterior timed phylogenies estimated from WGS data using a birth-death prior.

## 3 Results

### 3.1 Tree shapes capture transmission patterns

#### Different transmission networks result in quantitatively different tree shapes

To determine whether tree shapes captured information about the underlying disease transmission patterns within an outbreak, we simulated evolution of a bacterial genome over three types of outbreak contact network homogenous, super-spreading, and chain - and summarized the resulting phylogenies with five metrics describing tree shape. Figure 2 shows the distributions of these metrics across the three types of outbreaks, revealing clear differences in tree topology depending on the underlying host contact network. Super-spreader networks gave rise to phylogenies with higher Colless imbalance, longer ladder patterns, lower Δ*w*, and deeper trees than transmission networks with a homogeneous distribution of contacts. Trees derived from chain-like networks were less variable, deeper, more imbalanced, and narrower than the other trees. Other topological summary metrics considered did not resolve the three outbreak types as fully (Supplementary Information).

**Figure 2:**
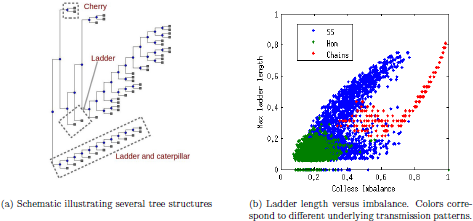
Distribution of simple summary measures of tree topology

**Figure 3:**
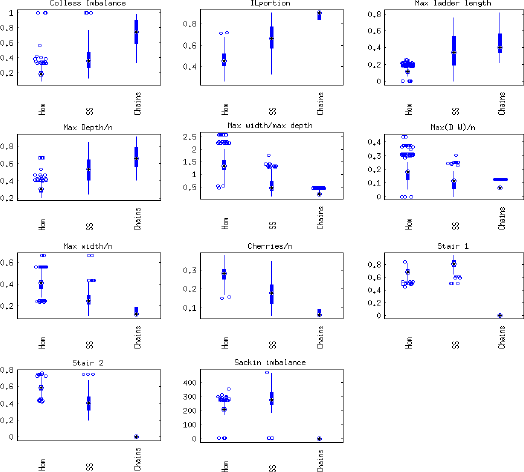
Box plots of the features used to summarize the shapes of phylogenies

### 3.2 Classification on the basis of tree shape

#### Topological metrics can be used to computationally classify outbreaks

To evaluate whether the five topological summary metrics could realiably and automatically differentiate between the three types of outbreaks, we trained a series of computational classifiers on the simulated datasets. We first trained a K-nearest-neighbour (KNN) classifier using the 11 tree features to discern which combinations of features correspond to phylogenies derived from the three underlying transmission processes. The KNN classifiers correctly identified the underlying transmission dynamics well (see Table 1 (a)), with an average of 89 (0.03)% of the homogeneous outbreaks, 86 (0.05)% of the super-spreader outbreaks and 100% of the chain outbreaks correctly classified under 10-fold cross-validation. Mis-classifications were between the homogeneous and super-spreader outbreaks.

**Table 1:** Results of cross-validated classification. Sensitivity (the true negative rate) here is the portion of homogeneous outbreaks correctly classified as homogeneous, and specificity (true positive rate) is the portion of super-spreader outbreaks correctly classified. For SVM classification, sensitivity and specificity have a trade-off, such that greater sensitivity can be achieved at the cost of reduced specificity and vice versa. Sensitivity and specificity are computed with the optimal threshold returned by matlab’s perfcurve function. The AUC captures the overall classifier quality. For K-nearest neighbour classification we report the portion correct by outbreak type, as there are three types. Numbers shown are mean (standard deviation) using the 10 classifiers found with 10-fold cross validation of the baseline case.

| KNN |  | Hom | SS | Chains |
| --- | --- | --- | --- | --- |
|  | Baseline | 0.89 (0.03) | 0.85 (0.05) | 1.00 (0) |
|  | Varied parameters | 0.90 (0.01) | 0.89 (0.01) | 1.00 (0) |
|  | Varied sampling | 0.98 (0.01) | 0.21 (0.01) | 1.00 (0) |
|  | Varied both | 0.98 (0.01) | 0.20 (0.01) | 1.00 (0) |
|  | 10 isolates | 0 (0.001) | 0.5 (0.5) | 0.5 (0.53) |
|  | 20 isolates | 0.99 (0.01) | 0.01 (0.001) | 0 (0) |
| SVM |  | Spec. | Sens. | AUC |
|  | Baseline | 0.93 (0.05) | 0.89 (0.05) | 0.98 (0.01) |
|  | Varied parameters | 0.92 (0.05) | 0.92 (0.04) | 0.98 (0.01) |
|  | Varied sampling | 0.79 (0.07) | 0.76 (0.05) | 0.86 (0.003) |
|  | Varied both | 0.83 (0.07) | 0.74 (0.06) | 0.87 (0.002) |
|  | 10 isolates | 1 (0) | 0 (0) | 0.61 (0.07) |
|  | 20 isolates | 0.93 (0.13) | 0.21 (0.34) | 0.78 (0.07) |

#### Support vector machine improves classification accuracy

To better resolve the separation between super-spreader-type outbreaks and those with homogeneous transmission, we trained a support vector machine (SVM) classifier to distinguish between those two types of outbreaks alone. Figure 4(a) shows the receiver-operator characteristic curve (ROC) for an SVM classification trained on 300 of the 1000 simulated homogeneous and super-spreader outbreaks. The area under the curve (AUC) is 0.97, reflecting a very good classifier performance; the theoretical maximum AUC is 1, and 0.5 corresponds to random guessing. We performed 10-fold cross-validation, each time training a new SVM on 900 of the 100 trees and testing it on the remaining 100. The average sensitivity was 0.93 and the average specificity was 0.89; the average AUC was 0.98. These values are listed in Table 1.

**Figure 4:**
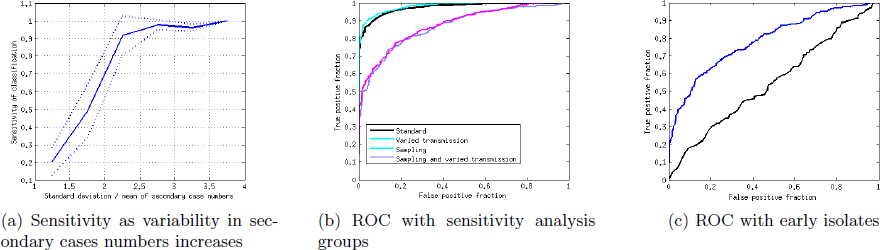
(a) Sensitivity of the SVM classification increases as the variability in the number of secondary cases in the outbreak increases. Variability is quantified as the ratio of the standard deviation to the mean of the numbers of secondary cases caused by an infectious case. Sensitivity is the portion of simulated outbreaks with the corresponding variability which were classed as super-spreader outbreaks; the solid line shows the mean sensitivity over the 10 SVMs produced by cross-validation and dotted lines are the mean *±* the standard deviation. (b, c) Receiver-operator characteristics for the SVM classifier based on the 11 summary metrics describing tree shape. ROC curves are a visual way to assess the classifier’s quality – a perfect classifier will obtain all the true positives and will have no false positives, giving an AUC of 1. An imperfect classifier has a trade-off, and can attain a specificity (true positive rate) of 1 at the cost of having a false positive rate of 1 (top right corner of the plot). The ROC curve illustrates the shape of this trade-off; the higher the area under the curve, the higher the quality of the classifier. Guessing yields an AUC of 0.5. In Figure 4(b), different lines correspond to the different groups of simulations in the SVM sensitivity analysis. Figure 4(c) shows the SVM classifier’s performance when only the earliest outbreak isolates are sampled. Performance is poor with 10 isolates (black line) and better with 20 (blue line).

### 3.3 Effects of the extent of super-spreading, sampling, and early classification

#### Outbreak classification is robust to variable parameters and model choice, but not to sampling

To explore how robustly phylogenetic structure captures variation in transmission processes, we performed sensitivity analyses in which we explored the effect of varying the transmission parameters *β/D*, sampling, and both the parameters and sampling together.

Using the KNN classifier applied to the three outbreak types, we found that the overall classifier error remained at approximately 10% when the transmission rate varied up to a factor of 2 (Table 1). The effect of reduced sampling density was much greater, and while the portion of homogeneous outbreaks correctly predicted was high (98%), the error was high because only 21% of super-spreader outbreaks were correctly classified. Mis-classification was between these two outbreak types, and chains of transmission were always correctly classified. Varying both the sampling and the parameters decreased the quality of the predictions slightly.

We also evaluated the sensitivity of SVM classification to different transmission model parameters by training and testing an SVM on a further 500 simulated super-spreading and homogenous networks, with variable transmission parameters *β/D*. As with the baseline parameter networks, the SVM returned an AUC of 0.98 for the variable *β/D* groups, though the sensitivity and specificity were both slightly reduced (0.92; see Table 1). However, the SVM’s performance declined with decreased sampling density (AUC of 0.86; sensitivity 0.76 and specificity 0.79), and decreased sampling with variable transmission parameters (AUC of 0.87, sensitivity 0.74 and specificity 0.83). Notably, the decline in performance was much less with the SVM method than the KNN method. Figure 4(b) shows the ROCs for SVM classification on these groups.

The decline in performance due to lower sampling density occurs for two primary reasons. The super-spreaders are relatively rare; if they were not, then the outbreak would not really be a ‘super-spreader’ outbreak, but one with a higher rate of transmission overall. When sampling density is reduced, there is therefore a good chance that the super-spreader individuals are not sampled. In addition, under weak sampling, only a few of a super-spreader’s secondary cases would be sampled. Both of these factors act to reduce the ability of the genealogy to capture super-spreading. Under very low sampling densities, it is likely that the probability of a given tree approaches what it would be under the homogeneous birth-death model or the appropriate coalescent model, even where the infectious period is not memory-less. Though we have not shown this, very low sampling should reduce the asymmetry that arises from one lineage continuing in the same host and another continuing in a new host, because each lineage would be expected to change hosts multiple times along a branch under low sampling densities. Accordingly, if sampling density is low enough that coalescent methods are appropriate, they may be used to relate branching times and some aspects of tree shapes to epidemic models [Frost and Volz, 2013].

We varied the extent of heterogeneity in the numbers of secondary cases in our outbreaks, using a negative binomial distribution and varying its parameters. We found that the classifier (trained on outbreaks each with a single superspreader but with varying secondary case numbers) had a high sensitivity of classification (*>* 0.7) when the ratio of the standard deviation to the mean of the secondary case number distribution was 2 or more. Figure 4(a) shows the average sensitivity increasing with the variability in secondary case numbers.

We tested the SVM classifiers to determine whether they could distinguish between phylogenetic trees derived from simulated sequence transmission on very different contact networks, namely dynamical models of sexual contact networks over a 5-year simulated time period [Robinson et al., 2013]. The performance was good when sampling was done over time, such that cases infected early in the simulation were likely to be sampled. When sampling was done at one time, years after seeding the simulated infection, neither classifier detected differences between the two types of contact network. Details are presented in the Supplementary Information.

#### Outbreak classification is possible using early isolates only

To determine whether classification of an outbreak is possible early in an outbreak-information that could potentially inform real-time deployment of a specific public health response-we evaluated the 10 KNN and 10 SVM classifiers’ performance when only the first 10 and first 20 genomes of the outbreak were sampled (10 of each, constructed using 10-fold cross-validation). The KNN performed poorly on the first 10 isolates, with none of the homogeneous outbreaks correctly classified, and only 50% of the others. Mis-classifications were between the super-spreader and chain outbreaks. After 20 isolates had been sampled, KNN classifiers grouped all outbreaks with homogeneous transmission. The SVM had AUC values of 0.61 and 0.78 after 10 and 20 isolates were detected, respectively (see Table 1 and Figure 4(c)), although the optimal cutoffs gave low sensitivity values. These data suggest that SVM classification can give some information about an outbreak’s transmission dynamics at early points within the outbreak.

### 3.4 Real-world outbreaks

#### Topological metric-based classification recapitulates known epidemiology of real-world outbreaks

Finally, to evaluate the classifiers’ performance on real-world outbreaks with known epidemiology, we applied the classifier to genome sequence data from two tuberculosis outbreaks whose underlying transmission dynamics have been described through comprehensive field and genomic epidemiology. Outbreak A [Gardy et al., 2011] was reported to arise from super-spreading activity, while Outbreak B displayed multiple waves of transmission, resulting in a somewhat more homogenous network.

We found that our classification results agreed with the empirical characterisations of the two outbreaks’ underlying transmission dynamics. In the KNN classification, Outbreak A was grouped with super-spreader outbreaks most often (56 (0.5) %), with 44% of the posterior trees grouping with homogeneous outbreaks none with chains. 77 (0.7) % of the trees from Outbreak B were classed as homogeneous, with the other 23% classed with super-spreader outbreaks. As above, numbers in parentheses are standard deviations over the 10 classifiers from the 10-fold cross-validation. The SVM classification grouped 75 (8) % of BEAST posterior Outbreak A trees with super-spreaders, and 76 (9) % of Outbreak B trees with homogeneous transmission. We also applied the classifiers to the maximum clade credibility (MCC) trees for the two outbreaks; the MCC tree from outbreak A grouped with super-spreaders and that from B grouped with homogeneous outbreaks in all of the 10 cross-validated classifications. Thus both classifiers’ predictions agree with epidemiological investigations of the outbreak, using tree shapes alone to classify transmission patterns.

## 4 Discussion

We have found that there are simple topological properties of phylogenetic trees which, when combined, are informative as to the underlying transmission patterns at work in an outbreak. Tree structures can be used as the basis of a classification system, able to describe an outbreak’s dynamics from genomic data alone. These topological signatures are robust to variation in the transmissibility, and to the nature and structure of the model, but sampling has a detrimental effect on the strength of the signal. Signs of the underlying transmission dynamics are present within the first 20 genomes sampled from an outbreak, and the classifiers are able to recapitulate known, real-world epidemiology from actual outbreak datasets.

The relationship between host contact heterogeneity and pathogen phylogenies is complex. In large datasets, phylogenetic branch lengths can reveal heterogeneous contact numbers [Stadler and Bonhoeffer, 2013], but distributions of branch lengths are not a suitable tool for small outbreaks of a chronically-infecting and slowly mutating organism like TB. Early work made the assumption that heterogeneous contact numbers would yield heterogeneous cluster sizes in viral phylogenies [Lewis et al., 2008]. But cluster sizes also depend on the pathogen population dynamics [Robinson et al., 2013] and the epidemic dynamics [Frost and Volz, 2013]. The relationship between heterogeneous contact numbers and tree imbalance [Leventhal et al., 2012] is not robust to the dynamics of a contact network [Robinson et al., 2013], sampling [Robinson et al., 2013, Frost and Volz, 2013] or the epidemic model used [Frost and Volz, 2013]. It is clear from this body of work that increased heterogeneity in contact numbers will not always lead to a simple increase or decrease of some measure, like imbalance, of tree structure. However, we have found that in small outbreaks, several simple topological features, taken together, can distinguish between outbreaks with high heterogeneity (a super-spreader) and low heterogeneity.

In any modelling endeavor, when a model reproduces features of real data – whether those are tree structures, branch lengths, or other data such as prevalence and incidence of an infection, locations of cases and so on – it remains possible that there are processes not included in the model that are the real origin of the observations. When we use models to interpret data, we use formal or informal priors to weigh the likelihoods of the assumptions behind the model compared to other processes that could drive the same phenomena. Here, one aspect of the complex relationship between contact heterogeneity and phylogeny structure is illustrated by the fact that genealogies from a long chain of transmission can look similar to genealogies derived from a super-spreader. Indeed, if one individual infects 10 others over a long period, and none of those infects anyone else, the genealogy among isolates would look the same as a genealogy in which each case infected precisely one other. However, it is very unlikely that such a chain of cases would occur, with no one *ever* infecting two others rather than one. Similarly, it is unlikely that one host could infect everyone in an outbreak, with no onward transmission by anyone else. In our simulations, once the occasional person in a long chain can infect two others, and if non-super-spreader individuals infect others homogeneously, we find that simple topological structures are well able to resolve the differences between chains and super-spreader outbreaks.

We have used 11 coarse and simple summaries of tree topology. However, any small set of a few summary statistics cannot capture the topology with much resolution. In contrast, most methods to compare phylogenies in fine detail are suited only for phylogenies on the same sets of tips [Robinson and Foulds, 1981], and so cannot be used to compare different outbreaks or to compare simulations to data. Finding the correct balance to summarize trees sufficiently that they can be compared across different tree sizes, different outbreaks and different settings, without summarizing them so much as to remove the most useful information is a challenge, and a number of methods will likely be developed, beginning with viral pathogens as in the recent work on Poon et al [Poon et al., 2013]. Indeed, while we feel that the measures we have used are demonstrative that tree structure is revealing, they are not intended to be comprehensive or exhaustive descriptions of tree topology. The fact that a few simple topological summaries can reveal underlying transmission patterns is a proof-of-principle that tree shape is informative.

We have taken a different approach than has recently been taken in a number of studies aiming to infer transmission trees from phylogenetic data [Jombart et al., 2014, Didelot et al., 2014, Jombart et al., 2010, Ypma et al., 2012, Ypma et al., 2013], or to identify or at least rule out transmission events based on epidemiological and genetic data [Köser et al., 2012, Walker et al., 2012, Grad et al., 2012, Török et al., 2013, Kato-Maeda et al., 2013]. These methods use the timing of case presentation (and estimated times of infection) to help determine who infected whom. In contrast, in pathogens with long and variable infectious duration, the timing of case presentation does not provide much information about the timing of infection. In this setting, even whole-genome sequence data may not contain sufficient information to clearly characterize individual transmission events, as we have recently found [Didelot et al., 2014]. However, individual transmission events are often of interest mainly because they reveal *patterns* of transmission. When we reconstruct an outbreak we are not seeking to determine whether case C will infect case D in the next outbreak, but rather, to find sufficient information about how the outbreak occurred that public health practices can benefit. Here we have found that tree shapes can reveal overall patterns of transmission without first inferring who infected whom.

The classification method we have developed provides not only an important empirical quantification of the degree to which genomic data is informative in the absence of epidemiological information, but is also a useful tool that can be used to describe outbreaks both retrospectively and prospectively. The ability to situate an outbreak on the spectrum from homogeneous transmission to super-spreading and to do so within the earliest stages of an outbreak when neither a large number of specimens nor detailed epidemiological information is available represents an important opportunity for public health investigations. Situating an outbreak on this spectrum does not require pinning down individual transmission events, but relies more on characterizing summary features of the outbreak and/or its phylogeny. If the data point towards a significant role for super-spreading in an outbreak, a containment strategy will require intensive screening of the super-spreader’s contacts. In an outbreak where onward transmission is occurring in chains, a focus on active case finding around multiple individuals will be needed instead. Ultimately, investigation of any outbreak of a communicable disease will involve the collation of multiple sources of information, including epidemiological, clinical, and genomic data. The approach described here represents one part of this toolbox, and has the advantages of being robust to the unique nature of complex chronic infection, providing useful information even when epidemiological information is incomplete, and being informative within the earliest stages of an outbreak.

## Acknowledgments

We thank the Vancouver Island Health Authority and the Interior Health Authority for their contributions to the real-world outbreak field investigations, and Dr. Patrick Tang, Dr. James Johnston, and Dr. Mabel Rodrigues for their contributions to the genome sequencing work. We thank Michael Boyd, who developed the phyloTop package with CC. CC is supported by the Engineering and Physical Sciences Research Council (EPSRC EP/K026003/1, EP/I031626/1).

