## Supplementary Information for "Phylogenetic tree shapes resolve disease transmission patterns" (from another version of this preprint: version 1, 2014-03-05)

### Robustness to the nature of the transmission model

We have trained classifiers using simple structural features of phylogenetic trees, and used classification to distinguish between outbreaks with a super-spreader, outbreaks with homogeneous transmission, and outbreaks built from chains of transmission. In the absence of real-world datasets in which the true transmission dynamics are known, we have used simulated outbreaks to train the classifiers. However, this necessitates specifying how the simulations are performed, and determining whether results are robust to our choice of model.

We applied our SVM classifier to simulated phylogenies described in Robinson et al [7]. In that work, dynamic networks of sexual contacts were created based on random graphs with a Poisson distribution (termed 'ER' because they are essentially Erdos-Renyi graphs), and with a distribution of contacts derived from the National Survey on Sexual Attitudes and Lifestyles (NATSAL) [3]. In our simulations, individuals with many contacts reported over the 5-year time frame in the survey had relationships of shorter durations, on average, than individuals who reported only a few contacts. In [7], when epidemics were sampled at one time after the epidemic had spread through the network for some years, the branch lengths, cluster sizes and Colless imbalances of phylogenies did not vary between NATSAL-type contact networks and ER-type networks. This is despite the fact that NATSAL-type networks have individuals with far more contacts than the mean (ie a "core group" of likely super-spreaders). However, when sampling was done over time, the phylogenies from NATSAL-type networks showed higher imbalance than those from ER-type networks, consistent with the work of Leventhal et al [4] on static networks. Here, we used the phylogenies from [7] with a duration of infectiousness $d = 40$ weeks, both under same-time sampling (homochronous) and sampling-through-time (heterochronous). Trees were inferred from simulated sequences using dnaml in the phylip package. Methods are described in detail in [7].

When we classified the homogeneous (ER) and super-spreader (NATSAL-derived) sexual contact networks, we found that the classification was poor on all groups of trees when hosts were sampled at the same time. Indeed, we found no differences between the distributions of any of the topological features we examined. However, in the heterochronous case, when individuals were sampled throughout the outbreak, the classifiers both performed very well. The KNN classifier classified 98.8% of the homogeneous-network trees correctly and all of the NATSAL-derived (super-spreader) trees correctly; the SVM classifier had no errors (AUC of 1), grouping NATSAL-derived trees with super-spreaders and ER-derived trees with homogeneous transmission. This is despite the fact that the parameters of the process, contact network, tree inference method and tree sizes were very different from the simulated homogenous, super-spreading, and chain networks the classifiers were trained upon. The basis of the sampling difference is that super-spreaders (or core group members) are likely to be infected early, and so are only included in the sample under heterochronous sampling. This result is consistent with [7], where only trees from heterochronous sampling showed differences in imbalance, branch lengths and cluster sizes between NATSAL and ER networks.

### Phylogenetic noise

In the results reported in the main text, we extract the true phylogeny from the simulations, based on the fact that we know who infected whom, and can therefore determine the time of the MRCA of all the nodes in the tree. In real outbreaks, this information is of course not available. To determine the extent to which our results are sensitive to phylogenetic noise, we created neighbour-joining trees from the simulations and applied the classification models to these.

We created neighbour-joining phylogenies as follows. The initial (seed) individuals were assigned a random sequence of A, C, T and G of 1000 base pairs in length. When an individual infected another individual, the transmitted sequence was mutated and the number of mutations reflected the time elapsed. As an example, suppose individual $i$ was infected with sequence $s_i$ at time $t_{0i}$ and then infected individuals $j$ and $k$ at times $t_{ij}$ and $t_{ik}$ with $t_{ij} < t_{ik}$. The time elapsed between $i$'s infection and $j$'s infection is $t_{ij} - t_{0i}$. Individual $j$ obtains a sequence $s_j$ which is $s_i$ with $m$ mutations, where $m$ is Poisson-distributed with mean $\mu(t_{ij} - t_{0i})$. Individual $k$ obtains a sequence $s_k$, which is

| Truth / Classification | Hom | SS | Ch |
|---|---|---|---|
| Hom | 704 | 302 | 6 |
| SS | 296 | 640 | 441 |
| Ch | 0 | 58 | 53 |

Table S1: KNN classification of neighbour-joining phylogenies

$s_j$ with a number of mutations distributed as Poiss($\mu(t_{ik} - t_{ij})$). Under this model, $k$'s sequence has the mutations occured in host $i$ before $i$ infected $j$, as well as further variation reflecting the fact that $k$ was infected at a point in time after $j$. This reflects the simplifying assumption, commonly made, that branching points in the phylogeny are very close to or the same as transmission events [10, 11, 9]. Sequences were sampled at the end of the duration of infection. The mutation rate $\mu$ was 6 SNPs per month - biologically unrealistic but required in the current scenario to generate sufficient genetic diversity over the short simulation timescales. If sufficient diversity is present for a phylogeny to be inferred, the mutation rate does not affect the structural properties computed here, and so does not affect the classification. Sampled simulated genome sequences from each outbreak case were aligned and phylogenetic trees for each of the networks were created using matlab's seqpdist and seqneighjoin functions [5]. Trees were constructed using a neighbour-joining method based on the Jukes-Cantor distance between sequences.

We found that phylogenetic noise has a large effect on the quality of KNN predictions and a modest effect on the SVM classification. It reduces the specificity and sensitivity of both classifiers, but SVM classification remains good. Table S1 shows the KNN results, illustrating that classification of chain-like patterns is particularly poor. Chain genealogies have a very characteristic shape, in which nearly all internal nodes of the tree have one tip descendant. Due to variable numbers of mutations between transmission events, the neighbour-joining trees fail to capture this structure sufficiently. Figure S1 shows the ROC when the SVM classifier is applied to homogeneous and super-spreader trees created with the neighbour-joining method.

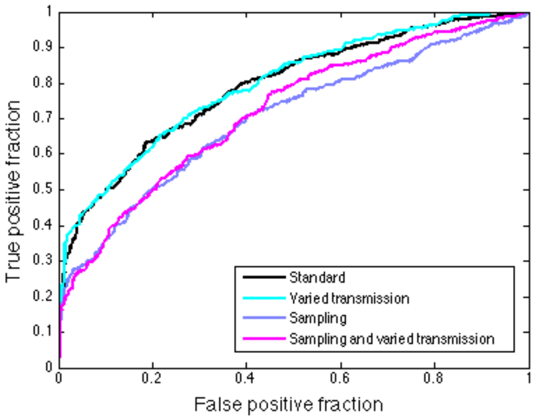


Figure S1: SVM classification (ROC curves) from the neighbour-joining phylogenies. Areas under the curve (AUCS) are 0.79, 0.79, 0.7 and 072 for the baseline, variable parameters, variable sampling and variable sampling with variable parameters lines respectively.

#### Additional topological summary measures

We considered a number of topological summary measures. These are listed in Table S2 and comprise different combinations of the ladder sizes, imbalance, depths, widths, and numbers of cherry formations [6] in the trees. Because they are not mutually independent, using more of these measures did not improve the classification.

Cherry formations, one of the features we examined, were studied in recent work by Frost and Volz [1]. They found expressions for the Sackin imbalance and number of cherries in virus phylodynamic models. A “cherry” is a pair of tips sharing a direct common ancestor, or equivalently, an internal node of the tree with two tip descendants [6]. Frost and Volz found that the number of cherries depended on the ratio of contact rates in two distinct subpopulations, and on the extent to which these subpopulations mixed preferentially within themselves. As such, it is not surprising that we did not find a strong signal in the cherry numbers, as their relationship to super-spreaders is complex. Also, our model is very different from that of Volz and Frost, in that we have modeled small outbreak trees in non-saturating outbreaks, and their work represents a viral pathogen (HIV) with large enough incidence and prevalence to be modeled with ordinary differential equations. Nonetheless, their approach gives an account of how and why the structures of topologies differ under different underlying epidemiological dynamics, including increased imbalance. Earlier work has also associated imbalance with positive selection and with super-spreaders [2, 4]; intuitively, a lineage corresponding to a super-spreader or one under positive selection would branch more rapidly than others, leaving asymmetric numbers of descendants on the right and left sides of descending sub-trees.

We have used what we term “ladders” in the trees; these are connected sets of internal nodes with a single leaf descendant. Our ladders are closely related to the “caterpillars” studied by Rosenberg [8]. The difference is that a caterpillar terminates in two tips (a cherry), whereas we include ladder formations further up in the tree, terminating in more complex sub-trees. See Figure S2 for an illustration. In our outbreaks, super-spreaders were infected early, and their sequences continued to mutate throughout the outbreak. As such, the signatures of super-spreading were likely to occur in the “ancestral” sequences – those of the super-spreader and their secondary infections prior to the mutations that occurred during the remainder of the outbreak. This motivates the use of structural features that occur internally in the tree, in contrast to cherries which reflect the dynamics among the most recently observed sequences. Cherry numbers would also be expected to change depending on sampling, though an exploration of how sampling affects phylogenetic structure is beyond the scope of this work.

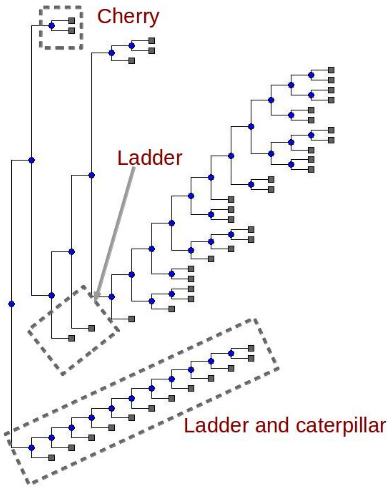


Figure S2: Illustration of a cherry formation, a ladder and a caterpillar in a phylogenetic tree.

### Model parameters

Figure S3 shows the distribution of infectious period in the model; this was the same for both types of networks. The association between topological structures and transmission via super-spreaders is dependent on these parameters, and in particular, when the duration of infection becomes quite long (greater than 2 years) and the value of $R$ decreases (below 1.3), the associations we report become weaker as a result of the very spread-out nature of transmission with these parameters. However, the results were robust to variation of the parameters such that $\beta/D$ was uniform in [1.25, 2.5].

| Name | Description | Relation to outbreak type |
|---|---|---|
| * Imbalance | Colless's $I$, defined in main text | Higher for super-spreader outbreaks |
| IL portion | Portion of internal nodes with one leaf descendant | No notable difference |
| * $\max(l/N)$ | Maximum ladder length / number of leaves | Higher for super-spreader outbreaks |
| IL nodes not in ladder | Number of internal nodes with one leaf descendant that are not in a ladder formation | No notable difference |
| * Scaled depth | Maximum depth / number of leaves | Higher for super-spreader outbreaks and anti-correlated with width/depth |
| * $\Delta w$ | As in main text; max difference in widths at successive depths | Lower for super-spreader outbreaks |
| * Maximum width/N | Maximum width over number of leaves | Lower for super-spreader outbreaks |
| Cherry number/N | Number of cherries over number of leaves | Slightly higher for super-spreader outbreaks |
| $\phi$ | Fraction of clades with very high imbalance | Higher for super-spreader outbreaks |
| Fraction high cherry numbers | Fraction of clades with very many cherries | Slightly lower for super-spreader outbreaks |

Table S2: Summary measures for phylogenetic trees. Those with an asterisk are the measures used to train the classifiers reported in the main text.

**Tree size**

While the structural features were normalised linearly, we did not ensure that they were scaled to be independent of tree size because the two real outbreaks were close in size. It is challenging to characterise the asymptotic expected value of structural features (see for example [8]). For outbreaks of the size reported here, asymptotic results may not be very meaningful in any case. For outbreaks of the size reported here, asymptotic results may not be very meaningful in any case. Furthermore, most of the work that has been done has focused on trees derived from one of several common simple models, including the Yule model. These may not be good descriptions of trees from densely sampled outbreaks, because in densely sampled outbreaks, the fact that infection processes are not memory-less affects the shape of the trees. When there is a new infection, the infectious period of the original host does not "reset", as would match the assumptions of homogeneous branching process models. Rather, the original host is likely to recover sooner than the newly-infected host, unless the infectious period is truly memory-less. For these reasons, we are uncertain how the structural features should scale on average with tree size, either in the Yule model or in a model adjusting for this asymmetry in the infectious period. We computed the structural features and the classification for trees from the homogeneous networks, and show the dependence on tree size in Figure S4. We found that the classification does depend on the size, which is likely one reason that classification performed poorly when only a few isolates had been observed. If we were interested in analysing much larger trees than those here, then it would be necessary to use summary features that capture more higher-resolution information about the tree at different scales than the very simple summary features we have used in any case. Simulation results such as the ones reported here could be the basis of conjectures as to how the structural summaries scale with tree size, but this analysis is beyond the scope of this work.

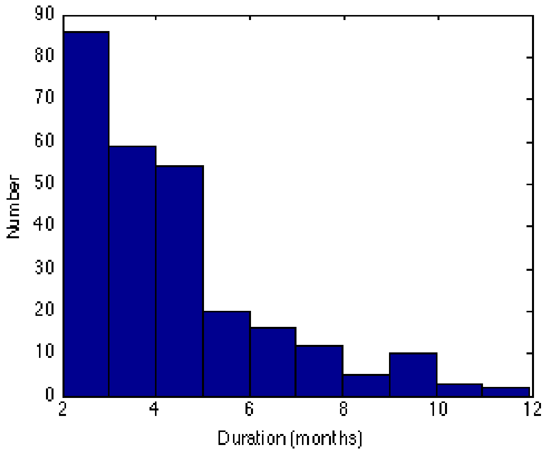


Figure S3: Distribution of lengths of infectious period in the transmission model

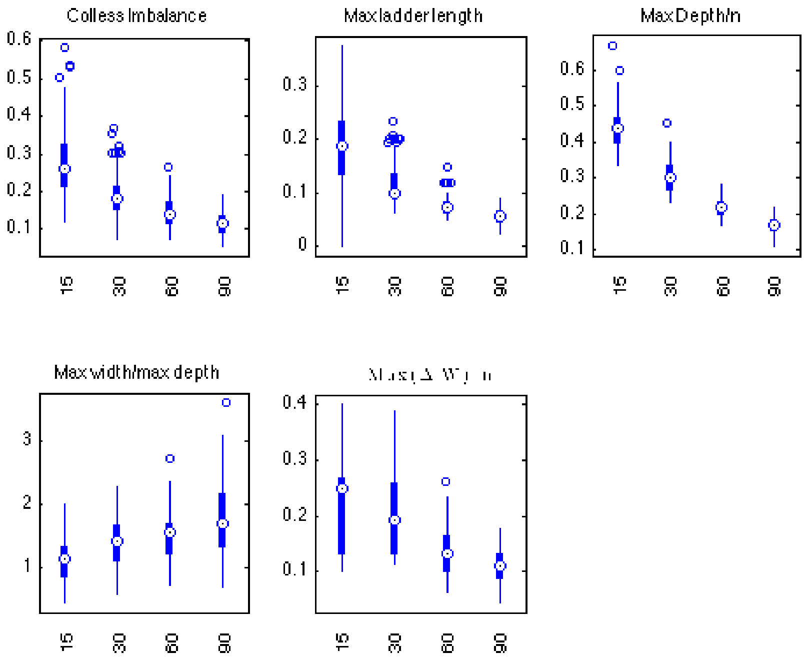


Figure S4: The structural features of trees, showing the dependence of the results on the number of tips included. Trees were derived from the homogeneous networks with the baseline parameter values reported in the main text.

[5] MATLAB. *version 7.14.0.739 (R2012a)*. The MathWorks Inc., Natick, Massachusetts, 2012.
